# Variation in virion phosphatidylserine content drives differential GAS6 binding among closely related flaviviruses

**DOI:** 10.1101/2025.08.09.669399

**Authors:** Lizhou Zhang, Byoung-Shik Shim, Claire Kitzmiller, Young-Chan Kwon, Audrey Stéphanie Richard, Michael Reynolds Farzan, Hyeryun Choe

**Affiliations:** Division of Infectious Disease, Boston Children’s Hospital, Boston, MA, USA; Department of Pediatrics, Harvard Medical School, Boston, MA, USA; Department of Immunology and Microbiology, UF-Scripps Biomedical Research, Jupiter, FL, USA

## Abstract

Many enveloped viruses engage phosphatidylserine (PS) receptors to enter cells, a phenomenon known as “apoptotic mimicry”. We previously reported that Zika virus (ZIKV), but not closely-related West Nile virus (WNV) or dengue virus (DENV), utilized AXL to infect cells because only ZIKV could bind the AXL ligand GAS6, a PS-binding protein. In this study, we investigated the mechanisms underlying the differential ability of these viruses to bind GAS6. Although immature virions expose larger patches of the viral membrane than do mature ones, our data show that virion maturity levels did not contribute to GAS6 binding. Surprisingly, while ZIKV contains PS comparable to cellular membranes, PS on WNV and DENV is markedly reduced. These findings explain why only ZIKV can bind GAS6 and provide insights into a novel mechanism by which closely-related flaviviruses differentially utilize cellular entry factors.

**IMPORTANCE:** Among flaviviruses, Zika virus uniquely causes microcephaly and congenital defects. While no flavivirus-specific entry receptors have been identified, they commonly take advantage of phosphatidylserine (PS) receptors to enter cells. Our previous studies revealed that Zika virus uniquely utilizes AXL, found in immune-privileged sites such as the brain and placenta, via binding to its ligand, GAS6. Our current study shows that despite being produced from the same cells, Zika virus has substantially higher PS content than closely related dengue and West Nile viruses, which likely explains Zika virus’s unique ability to bind GAS6. These findings provide insight into how closely related flaviviruses can vary substantially in their use of cellular entry factors, potentially contributing to the distinct diseases they cause.

## INTRODUCTION

The genus *Flavivirus* comprises enveloped viruses that include important human pathogens such as Zika virus (ZIKV), West Nile virus (WNV), dengue virus (DENV), and yellow-fever virus (YFV), as well as Japanese encephalitis virus and tick-born encephalitis virus. These viruses are transmitted through the bites of infected arthropods, such as mosquitos or ticks. For many of them, humans are dead-end hosts because they cannot amplify the virus to a sufficient titer to reinfect insect vectors. Exceptions to this include YFV, DENV and ZIKV, which caused multiple repeated global epidemics and have become endemic to many geographical regions. Although these viruses share similarities in sequence and structure, ZIKV uniquely causes congenital infections leading to microcephaly and other birth defects. In contrast, WNV predominantly infects the central nervous system, causing inflammatory diseases mainly in adults (1, 2) but is rarely associated with microcephaly (3, 4). Neither DENV nor YFV are neurotropic and have not been linked to microcephaly. Extensive evidence from human and animal studies (5–12) support a causal relationship between ZIKV and microcephaly, distinguishing it from other flaviviruses.

Flaviviruses replicate in the cytoplasm. Their ∼11kb genome is translated as a single polypeptide and cleaved by cellular and viral proteases into the structural and non-structural proteins. The structural proteins include capsid (C), pre-membrane (prM), and envelope (E), and the non-structural proteins consist of NS1, NS2A, NS2B, NS3, NS4A, NS4B, and NS5. Flavivirus virions assemble on the cytoplasmic side of the expanded endoplasmic reticulum (ER), bud into the ER lumen, and undergo maturation as they transit through the ER and Golgi apparatus. During virion maturation, the prM protein, which is essential for E protein folding and assembly (13), is cleaved into the pr peptide and M protein by the cellular protease furin in the Golgi apparatus (14). The pr peptide remains associated with the virion within the cell but is released when the virion reaches neutral pH of the extracellular environment (15). Immature virions have a spiky surface because E proteins that are associated with the uncleaved prM proteins stand upright as trimers on the virion surface. In contrast, mature virions are smooth because after the cleaved pr peptides are released, E-proteins form dimers that lie flat on the virion surface. There are 90 such E-protein dimers per virion, and each E protein is paired with an M protein underneath (16). This arrangement completely covers the virion, leaving minimal membrane exposure. However, the dynamic “breathing” motion of viral proteins and membranes (17, 18) transiently exposes patches of membrane, particularly at 37°C (19). Compared to mature virions, immature virions expose more of their membrane due to the upright configuration of prM-E trimer spikes (20). In addition to fully mature and immature virions, chimeric or mosaic forms, which are partly mature and immature in patches, have been observed by electron microscopy (21, 22). While the mature state is generally required for infectivity, immature virions can still become infectious if pr peptide cleavage occurs by target cell furin during infection (23). Moreover, mosaic virions are infectious even without target-cell furin (24).

Virus entry into the target cell is typically initiated by the viral entry glycoprotein binding to a receptor expressed on the target-cell surface. In the case of flaviviruses, no virus-specific entry receptor has been identified, but many cell-surface molecules are known to facilitate their infection. Best known entry factors include C-type lectins (namely, DC-SIGN/CD209 and L-SIGN/CD299) and various PS receptors (25–33). Lectins recognize carbohydrate moieties on the virion surface, while PS receptors bind PS present in the virion membrane (27, 28). The primary physiological function of PS receptors is to bind the PS exposed on the apoptotic cells and mediates their phagocytic clearance, a process known as efferocytosis (34–38). This process avoids immune activation to prevent responses against self-proteins. Thus, many enveloped viruses exploit these PS receptors to gain entry into cells, and this strategy is known as “apoptotic mimicry” (39, 40). Among PS receptors, the T-cell immunoglobulin mucin (TIM) family (TIM1, TIM3 and TIM4) and the TAM family (TYRO3, AXL, and MERTK) receptors play key roles in flavivirus infection (26–28, 31–33, 40). Of the three human TIM-family members (TIM1, 3 and 4), TIM4 is expressed on professional phagocytes such as dendritic cells and macrophage sub-populations (36, 41). In contrast, TIM1 is induced to express on non-professional phagocytes, namely, epithelial cells in tissues such as the lung, kidney, mammary gland, and placenta, where it clears neighboring cells when they undergo apoptosis (42–50). The TAM-family members are broadly expressed across various tissues, particularly in regions subject to continuous challenge and renewal. They are expressed in the endothelium and immune privileged sites such as the brain, eye, placenta, and testis, where they facilitate apoptotic cell clearance while maintaining the integrity of vasculature and barrier cells of those sensitive areas (9, 47, 51–55).

While TIM-family receptors directly bind PS, TAM family receptors engage PS indirectly through soluble mediators, specifically, growth arrest specific gene 6 (GAS6) and Protein S (ProS), which are present in serum and other bodily fluids (56, 57). GAS6 serves as a ligand for all three TAM receptors, whereas ProS binds only to TYRO3 and MERTK (57). Consequently, GAS6 is the sole ligand for AXL. Although PS is the primary target for these PS-binding receptors, a subset of them, such as human TIM1, MFGE8, Annexin A5 also bind phosphatidylethanolamine (PE) with equal or greater efficiency compared to PS (28, 58–60). Like PS, PE is typically restricted to the cytoplasmic leaflet of the membrane lipid bilayer and flips to the extracellular side during apoptosis (61), which also facilitates efferocytosis and virus entry (28).

GAS6/AXL-mediated TAM receptor usage by enveloped viruses was reported for the first time by Morizono et al. (32) using lentiviral vectors pseudotyped with alphavirus glycoproteins derived from Sindbis or Ross River virus. We also made similar observations: retroviral vectors pseudotyped with glycoproteins from filoviruses, arenaviruses, or alphaviruses showed markedly enhanced entry mediated by AXL (26). Surprisingly, in the same study, we found that WNV virus-like particle (VLP) did not use AXL, although it efficiently used TIM1 (26). In fact, among all TIM1-using viruses, WNV VLP was the only one that failed to utilize AXL. This was the first time we noticed WNV’s inability to engage AXL. Later, we observed that DENV, like WNV, also failed to use AXL, and demonstrated that such differential use of AXL was due to ZIKV’s unique ability to bind GAS6, while WNV and DENV do not (31). The mechanism behind this intriguing observation remains unclear. In the current study, we investigated the differential ability of these viruses to bind GAS6 and found that WNV and DENV contain significantly lower PS levels in their virions. This low PS content in WNV and DENV virions does not support their interaction with GAS6, whereas the higher PS content in ZIKV facilitates it.

## RESULTS

### ZIKV, but not WNV or DENV, binds GAS6

We have previously reported that ZIKV, but not WNV or DENV, efficiently uses AXL and that such differential AXL use is due to the unique ability of ZIKV to bind GAS6: e.g., WNV and DENV do not bind GAS6 (31). However, the underlying mechanism remains unknown. To investigate the mechanism behind this unexpected phenomenon, we first confirm the reproducibility of our previous observations using viruses produced in human cell lines, because in our previous study, the viruses were grown in the monkey cell line, Vero (31). In this study, viruses were propagated in A549, a human lung epithelial cell line, or in Huh7, a human hepatic epithelial cell line. To measure GAS6 binding, these viruses were incubated with the Ig-fusion form of GAS6 (GAS6-Ig), and captured virus particles were precipitated, using Protein A-Sepharose beads. These captured viruses were then visualized by Western blots using anti-E protein antibodies or quantified by RT-qPCR of viral RNA. Ig-fusion forms of C-terminal half of GAS6 (C-GAS6-Ig), which binds AXL but does not bind PS, and TIM1-Ig, which binds all three viruses, were included as a negative control and positive control, respectively (28, 31). The reason TIM1-Ig could bind all three viruses, although it is a PS-binding protein, like GAS6, is because it is able to bind PE in addition to PS and because the PE levels are typically much higher than those of PS (28). Figure 1 shows that ZIKV, but not WNV or DENV, bound GAS6-Ig as shown both by WB (Figure 1A) and RT-qPCR (Figure 1B). The results obtained from the viruses produced in A549 and Huh7 are nearly identical. As expected, C-GAS6-Ig did not bind any virus, and TIM1-Ig bound all viruses. These data confirm, using live viruses produced from human cell lines, our previous report that ZIKV, but not WNV or DENV, can bind GAS6 and utilizes AXL as an entry factor (31).

**Figure 1.**
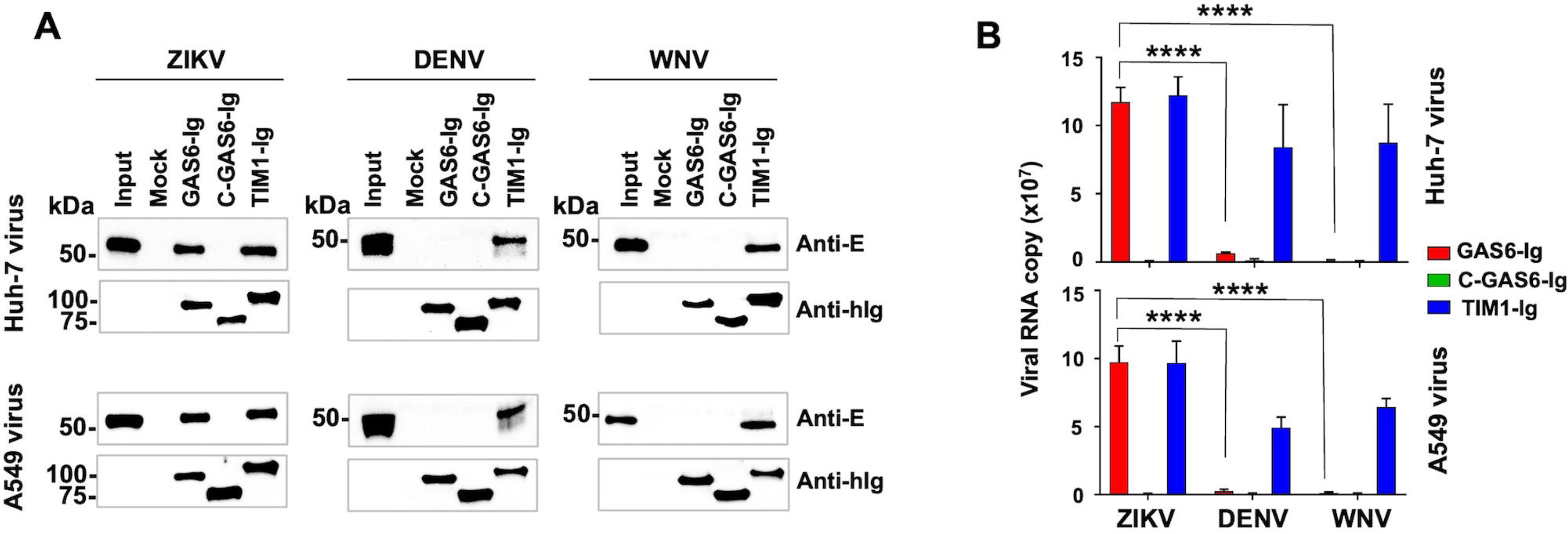
ZIKV, but not WNV or DENV, binds GAS6. ZIKV, DENV, or WNV produced from Huh-7 or A549 cells were incubated with the indicated Ig-fusion protein or Mock sup and precipitated using Protein A-Sepharose beads. **(A)** Bound viruses were analyzed by non-reducing (E proteins) and reducing (Ig-fusion proteins) SDS-PAGE and WB. ZIKV and DENV E proteins were detected by 4G2 antibody and WNV E protein by 3.91D antibody. Anti-human IgG antibody was used to detect the Ig-fusion proteins. **(B)** Bound viruses were quantified by RT-qPCR using the primers and probe targeting the NS1 gene of each virus and whose sequences are listed in Table 1. Data are represented as the mean ± SD of three independent experiments. Non-specific virus binding in the control, Mock sup, is subtracted from the values of each experimental sample. Statistical significance of the difference between ZIKV vs DENV and WNV was analyzed by one-way ANOVA (\*\*\*\**p* < 0.0001).

### Virion maturity does not alter GAS6 binding

To explore the mechanism with which closely-related flaviviruses differentially bind GAS6, we first focused on virion maturity, because immature virions expose more membrane than do mature viruses, owing to different E protein arrangement on the virion surface (20). We therefore reduced WNV virion maturity to promote binding of GAS6 by immature WNV. To do so, we interfered with WNV maturation by treating infected cells with a furin inhibitor, Decanoyl-RVKR-CMK. This treatment reduces virion maturation by inhibiting the cleavage between the pr peptide and M protein. Vero cells treated with Decanoyl-RVKR-CMK produced an immature virion population, as indicated by the uncleaved prM band, while untreated Vero cells produced mature WNV virions (Figure 2A, left panel). WNV so treated still did not bind GAS6-Ig (Figure 2A, right panel), while it bound TIM1-Ig. We then tried an alternative method to produce immature virions. Based on our previous experience, flaviviruses grown at lower temperature exhibit lower maturity. Accordingly, cells infected with WNV were cultured at 37°C or 28°C. As expected, WNV virions produced at 28°C were less mature than those produced at 37°C, demonstrated by the uncleaved prM band detected only in the WNV grown at 28°C (Figure 2B, left panel). However, this 28°C-produced WNV again did not bind GAS6-Ig, while it did bind TIM1-Ig (Figure 2B, right panel). Finally, assuming the WNV produced under these two conditions may not be sufficiently immature, we produced completely immature WNV from LoVo cells. This human colon carcinoma cell line lacks functional furin due to a frame-shift mutation (62). WNV produced from LoVo cells was therefore fully immature, as indicated by absence of the cleaved M protein (Figure 2C, left panel). Nonetheless, this completely immature WNV still did not bind GAS6-Ig (Figure 2C, right panel). These data demonstrate that virion maturity is unlikely to contribute to GAS6 binding.

**Figure 2.**
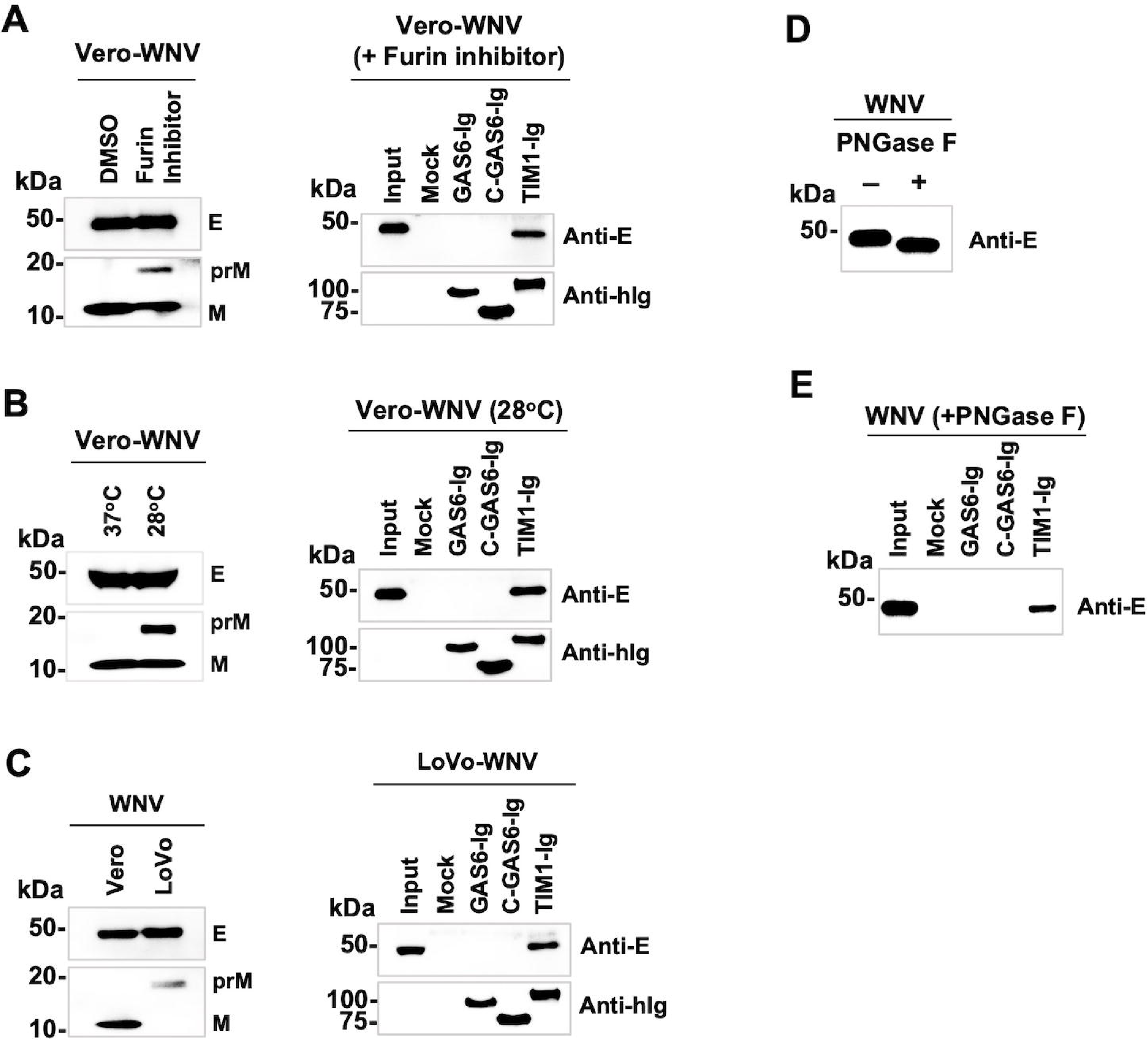
Virion maturity and N-glycans do not alter GAS6 binding. WNV grown in Vero or LoVo cells with or without the indicated treatment was incubated with the Ig-fusion protein or Mock sup and precipitated using Protein A-Sepharose beads and analyzed by WB. 3.91D antibody was used to detect the E protein, an anti-WNV M antibody to detect the M and prM proteins, and an anti-human IgG antibody to detect the Ig-fusion proteins. **(A)** WNV was grown in Vero cells at 37°C in the presence or absence of a furin inhibitor, Decanoyl-RVKR-CMK, at 50 uM. **(B)** Vero cells infected with WNV were cultured at 37°C or 28°C, and virus was harvested 2d later. **(C)** WNV was grown in Vero or LoVo cells at 37°C and harvested 2d (Vero cells) or 3d (LoVo cells) later. (A-C) Shown are representative results from two independent experiments. **(D)** 50 ul WNV produced in Vero cells was treated with or without PNGase F at 20 units/μl and analyzed by WB. PNGase F treated E protein migrated faster, indicating removal of N-glycans. **(E)** The same viruses in A were incubated with the indicated Ig-fusion proteins, precipitated with Protein A-Sepharose beads, and analyzed by SDS-PAGE and WB. E protein was detected using 3.91D antibody.

### N-glycans on the E protein do not affect GAS6 binding

We next investigated whether N-linked glycans on the virion contribute to GAS6 binding. N-glycans on viral proteins, including flavivirus E proteins, play more than just a shielding role against immunity. They have been linked to enhanced virus production in mosquito or avian hosts, tissue tropism, and an increased ability for neuroinvasion (63–66). Lack of N-glycosylation has also been linked to attenuated phenotype or reduced virulence (67). Although GAS6 has not so far been shown to directly bind carbohydrates, we nonetheless sought to investigate N-glycans’ contribution to differential GAS6 binding because N-glycans on the virion surface could sterically hinder GAS6 binding to the virion membrane. Therefore, to study their effect, N-linked glycans on the virions were removed by incubating WNV with PNGase F. As Figure 2D shows, PNGase F treatment reduced the apparent size of WNV E protein as expected from the removal of N-glycans. However, PNGase F treatment did not make WNV bind GAS6-Ig (Figure 2E), indicating that the different number of N-glycans on the three viruses does not account for differential GAS6 binding by these viruses.

### Virion phospholipids, not viral proteins, mediate ZIKV binding to GAS6

To confirm that capturing ZIKV by GAS6-Ig in our assays is mediated by PS, not by proteins, on the virion, we treated the ZIKV-GAS6-Ig complex with phospholipase C (PLC) after virions are captured by GAS6-Ig. PLC cleaves the bond between phosphate and glycerol in all phospholipids, removing their headgroup from the diacylglycerol backbone (Figure 3A). Therefore, if ZIKV binds GAS6 through PS, PLC treatment should dissociate the virions from GAS6-Ig (Figure 3B), but if ZIKV-GAS6 binding occurs through viral proteins, PLC treatment will not make any difference. As shown in Figure 3C (left panel vs right panel), PLC digestion removed a majority of ZIKV bound to GAS6 or TIM1. Similarly, pre-treating ZIKV with PLC before its binding to GAS6-Ig or TIM1-Ig yielded similar results (Figure 3C, middle panel). Virus binding to TIM1 is reduced or eliminated by PLC digestion, because PLC cleaves the head group of all phospholipids. Quantification by RT-qPCR of captured ZIKV digested by PLC over a range of concentrations produced similar results (Figure 3D). These findings confirm that ZIKV binding to GAS6 is mediated by phospholipids rather than viral proteins.

**Figure 3.**
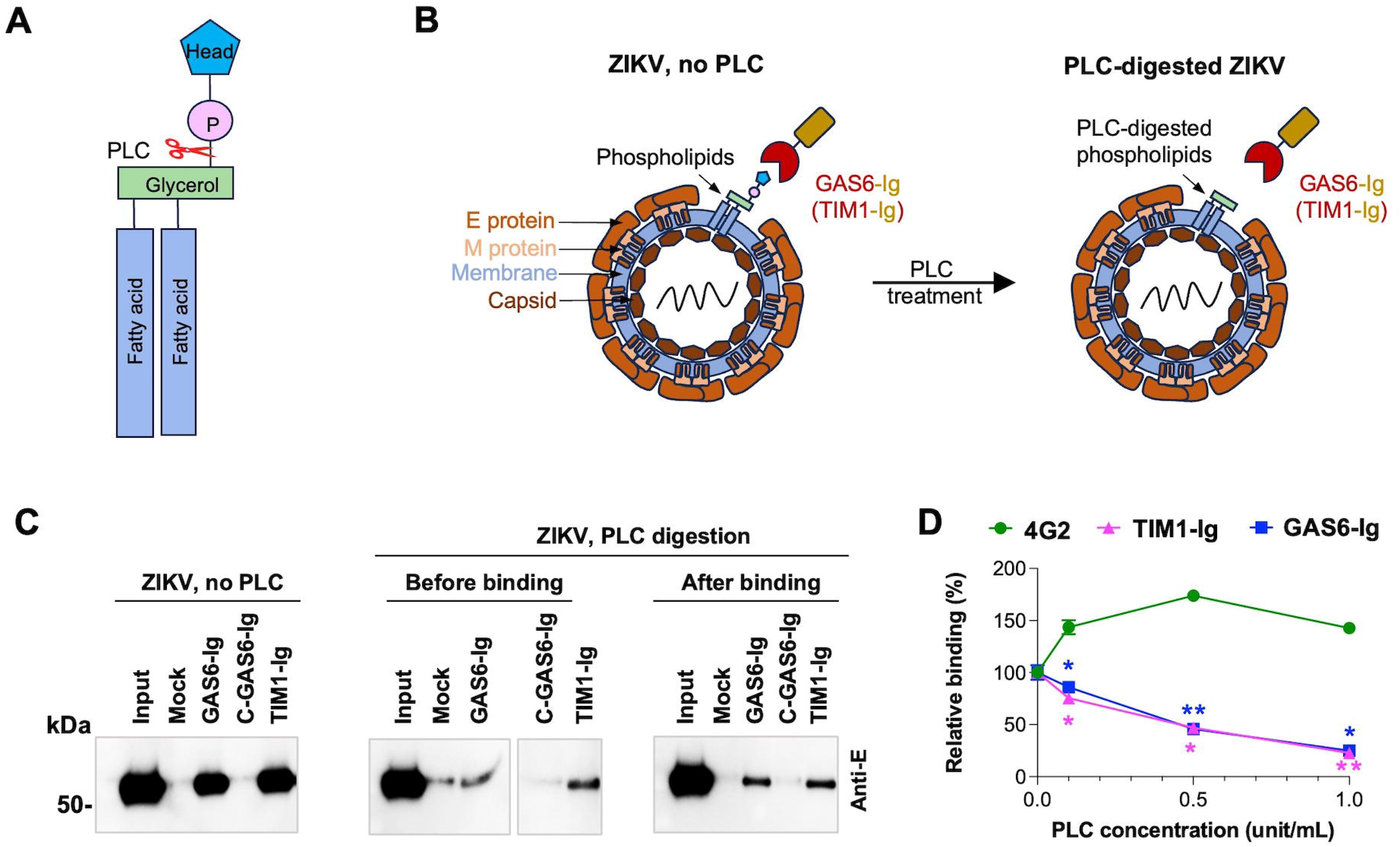
Virion phospholipids, not viral proteins, mediate ZIKV binding to GAS6. **(A)** A diagram of a phospholipid molecule with the cleavage site for phospholipases C (PLC) indicated. **(B)** A schematic presentation of GAS6-Ig or TIM1-Ig binding to PS in the ZIKV virion with or without PLC digestion. **(C)** ZIKV grown in Vero cells were incubated with the indicated Ig-fusion proteins, precipitated with Protein A-Sepharose beads, and analyzed by WB, using 4G2 antibody. ZIKV was either digested or not digested with PLC before or after incubation with the Ig-fusion proteins as indicated. (**D**) ZIKV particles were either treated or not treated with increasing concentrations of PLC, then precipitated with 4G2 antibody, TIM1-Ig, or GAS-Ig, and analyzed by RT-qPCR to detect the viral NS1 gene. Relative binding was calculated by normalizing the PLC-treated ZIKV to the PLC non-treated one. Statistical significance of the TIM1-Ig or GAS6-Ig group to the 4G2 group at the same PLC concentration was analyzed using repeated measures ANOVA (\**p* < 0.05; \*\**p* < 0.01) and is presented as the mean ± SD.

### PS content of ZIKV is substantially higher than that of WNV and DENV

Having confirmed that ZIKV binds GAS6 via phospholipids in the virion membrane, we next investigated whether the phospholipid content of DENV, WNV, and ZIKV is different. Initially, we did not pursue this hypothesis because it seemed the least likely because the membranes of enveloped viruses are generally believed to derive from cellular membranes. In addition, all three viruses were grown in the same cell line, and all of them bud from the same organelle, the ER. Nevertheless, in the absence of a clear mechanistic explanation, we compared the phospholipid contents of purified ZIKV, WNV and DENV grown in Vero cells, using two-dimensional thin layer chromatography (2D TLC). Before analyzing viral phospholipids, we verified the positions of the four major phospholipids—phosphatidylcholine (PC), PE, PS, and phosphatidylinositol (PI)— on the 2D TLC plate. Although sphingomyelin (SPH) is not a phospholipid, it contains a phosphate group and thus is stainable by iodine, so it was included in the assay. These lipids were purchased from Avanti Polar Lipids, mixed in a ratio (PC:PE:PS:SPH:PI=10:5:2:2:1) roughly comparable to that of mammalian cell membrane (68–70), separated by 2D TLC on a silica plate, and stained with iodine vapor. The location of each of these phospholipids is indicated in the upper panel of Figure 4A. We then analyzed the lipids extracted from the ER membrane of Vero cells in the same way and compared the results to those of the commercial phospholipids. The five lipids derived from Vero ER membrane are located at the same positions as the commercial ones (Figure 4A, lower panel). Commercial lipids formed more compact spots compared to those from Vero ER membrane, as they contain homogenous fatty acyl chains (we purchased those with dioleic acid), whereas the fatty acyl chains of cellular lipids are heterogenous in length.

**Figure 4.**
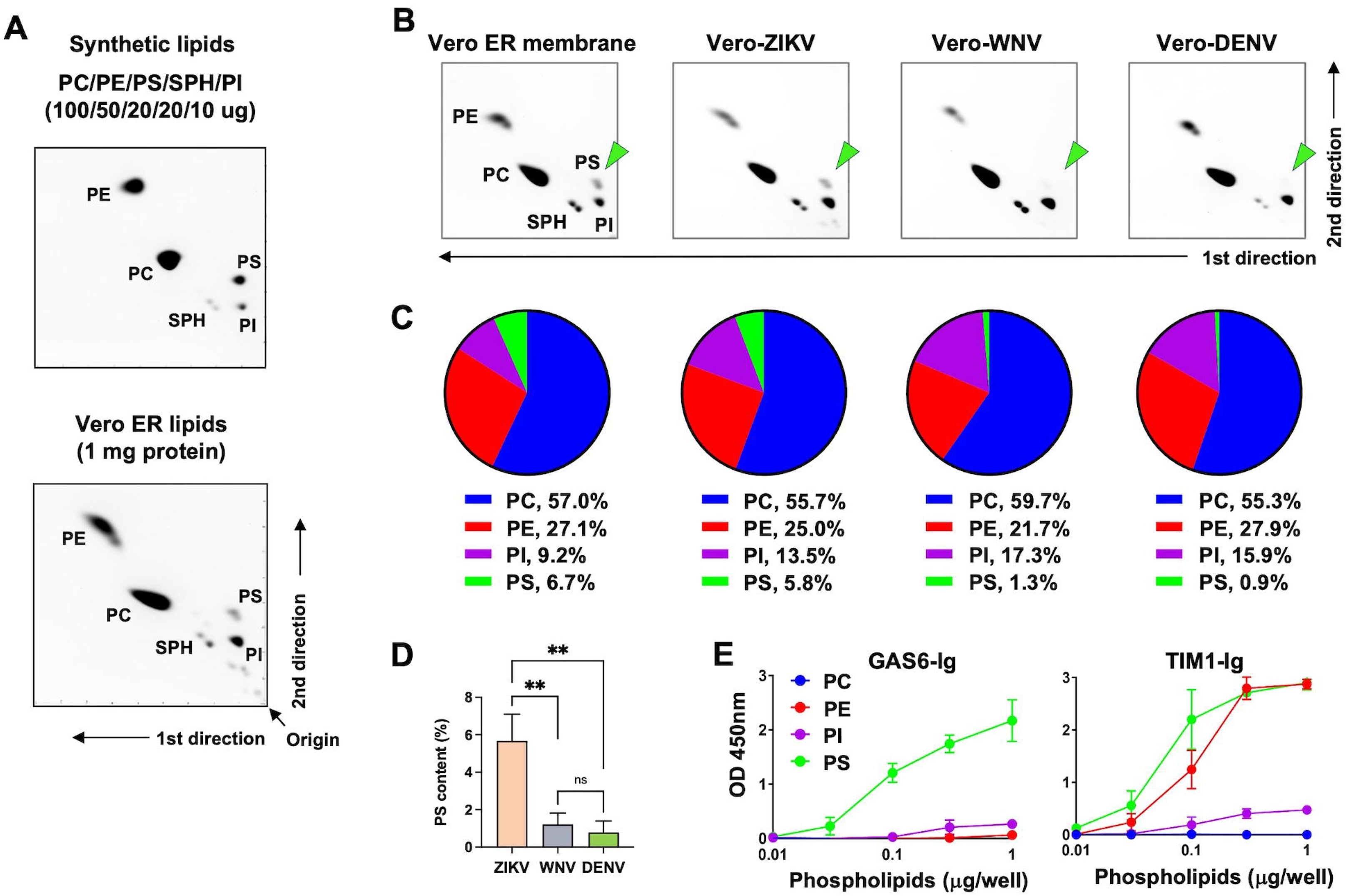
PS content of ZIKV is substantially higher than that of WNV and DENV. **(A)** Two-dimensional thin layer chromatography (2D TLC) of synthetic and Vero ER phospholipids. Upper panel: Synthetic DOPC, DOPE, DOPS, DOPI and SPH were mixed at the indicated amount, separated by 2D TLC, and stained with iodine vapor. The location of each lipid is marked. Lower panel: Total lipids were extracted from the ER membrane isolated from Vero cells, using the method of Bligh and Dyer, and analyzed and stained in the same way as the synthetic lipids. **(B**) Total lipids were either extracted from the ER membrane isolated from Vero cells or from the indicated virus grown in Vero cells and purified using potassium tartrate gradient centrifugation. Extracted lipids were separated by 2D TLC, and phospholipids were visualized by iodine vapor. Each image is a representative of three independent experiments, conducted using three independent sample preparations. All three experiments are shown in Figure S2. **(C)** Phospholipid composition was determined by densitometry from four consecutive images of each plate, taken every two minutes, and averaging across three independent experiments. The resulting average values from these experiments are represented as pie charts, with the numerical values shown underneath. **(D)** Statistical significance of the difference in PS content among ZIKV, DENV, and WNV shown in C was analyzed by one-way ANOVA (\*\**p* < 0.01) and is presented as the mean ± SD of twelve images, with four images from each of three independent experiments. **(E)** Phospholipid ELISA assays were conducted by coating polystyrene plates with increasing amount of the indicated phospholipid and incubating them with 3 nM GAS6-Ig or TIM1-Ig. Shown are representatives from three independent experiments with similar results.

Having analyzed phospholipids of Vero ER membrane, we then analyzed the lipids from purified ZIKV, WNV, and DENV. These viruses were propagated in Vero cells and purified using potassium tartrate gradient centrifugation. Their purity was assessed by Coomassie blue staining of viral proteins after they were separated by electrophoresis. Figure S1 shows these virus preparations were highly pure. Total lipids were extracted from these viruses and analyzed by 2D TLC (Figure 4B). One immediately noticeable difference is that the PS spots of WNV and DENV are barely visible, while those of ZIKV and Vero ER membrane are clearly visible with comparable intensity. When quantified (Figure 4C and Figure S2), the PS content of ZIKV (5.8%) was comparable to that of Vero ER (6.7%), while the PS contents of WNV and DENV were much lower (1.3% and 0.9%, respectively). ZIKV has significantly higher PS content (4.5-6.4 folds) compared to WNV and DENV (Figure 4D). PC and PE contents of all viruses were roughly comparable to those of Vero ER, although PE of WNV (22%) is modestly lower than that of ZIKV and DENV (25-28%) and Vero ER (27%). Interestingly, as Figure 4C shows, the PI level of all three viruses was considerably higher (13.5-17.3%) than that of the Vero ER membrane (9.2%). However, since it increased in all three viruses, PI cannot be the factor responsible for the differential GAS6 binding by the viruses.

Because PI is also negatively charged like PS and because a subset of PS-binding proteins, such as TIM1 and MFGE8, bind additional phospholipid PE (28, 58), we nonetheless analyzed the binding profile of GAS6 to each phospholipid to verify that PS, but not other phospholipids, is responsible for GAS6 binding. We conducted lipid ELISA with PC, PE, PS, and PI. As Figure 4E shows, GAS6 binds only PS, while TIM1 binds both PE and PS as we reported previously (31). Although PI weakly binds GAS6 and TIM1 at high concentrations, it cannot explain why WNV and DENV do not bind GAS6, as PI is modestly more abundant in WNV and DENV than in ZIKV (Figure 4C). These data show that the PS content of ZIKV is higher than that of WNV or DENV and correlates with ZIKV’s ability to bind GAS6.

### PL composition of the ER membrane is not differentially altered by different viruses

To further investigate the mechanism with which ZIKV, WNV, and DENV differentially incorporate PS into their virion, we first examined the changes in the ER lipidome induced by virus infection. The ER lipids were extracted when the majority of cells were infected, which was two days after infection for WNV and ZIKV and six days for DENV. The viruses used in this study were also harvested two days (WNV and ZIKV) or six days (DENV) after infection. The ER lipids from the infected Vero cells were compared to those extracted from uninfected cells. As Figure 5A, and Figure S3 show, no substantial alteration in the phospholipid composition of the ER membrane was induced by virus infection or by different viruses. Moderately higher PS level was observed in the cells infected with WNV or DENV, compared to uninfected cells or those infected with ZIKV (Figure 5B and 5C), which cannot explain the low PS content observed with WNV and DENV. These results demonstrate that virus infection does not significantly alter the ER lipidome, at least for phospholipid ratio, and thus cannot explain the apparent difference in PS content observed among the three viruses. These data also suggest that viruses themselves are likely responsible for differential PS incorporation into their virion.

**Figure 5.**
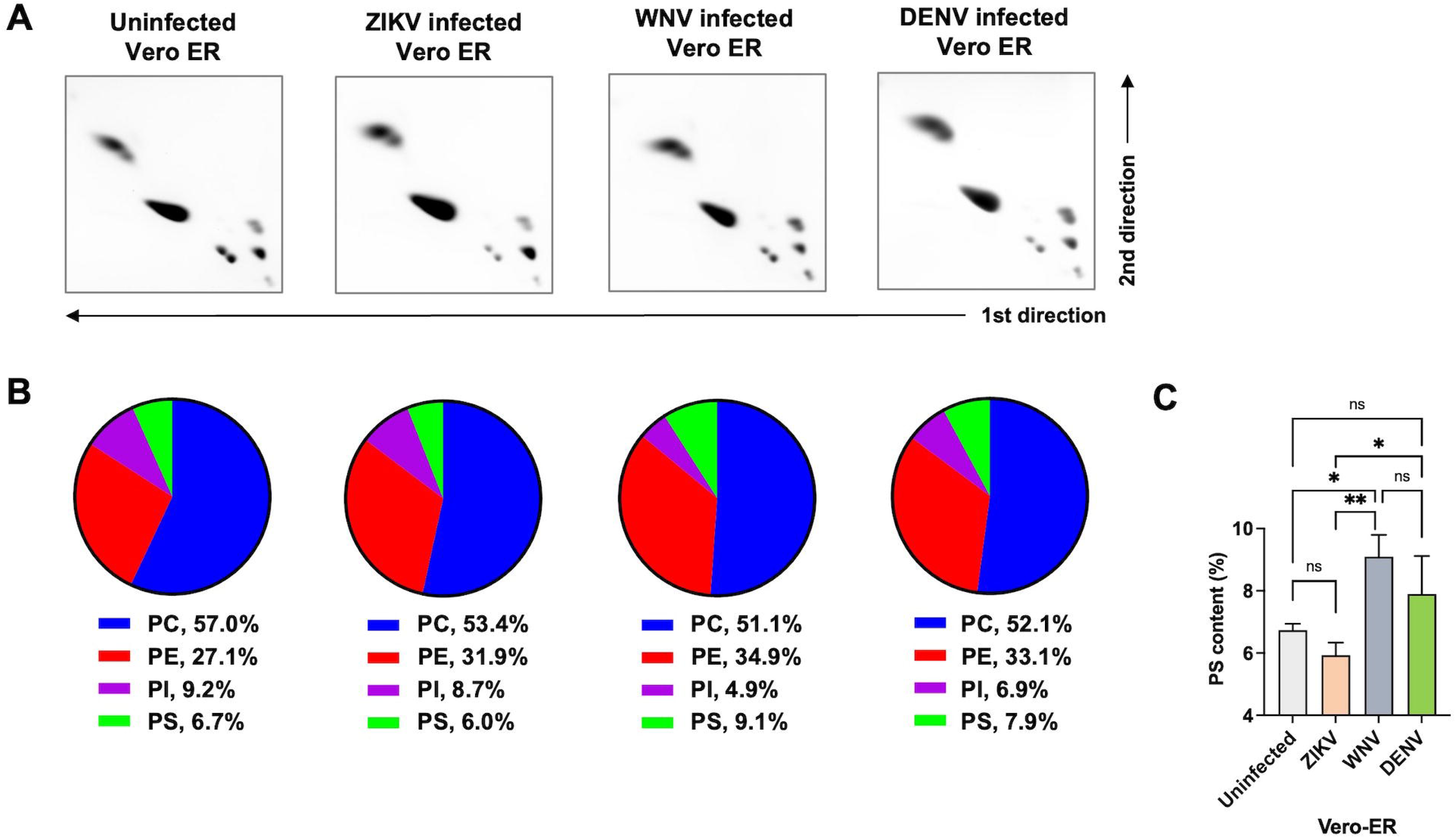
Phospholipid composition of the ER membrane is not differentially altered by different viruses. **(A)** Vero cells were uninfected or infected with ZIKV, WNV or DENV at an MOI of 1, and the ER membranes were extracted from these cells 2 days (ZIKV or WNV infected) or 6 days (DENV infected) later. Total lipids were extracted and separated by 2D TLC, and phospholipids were visualized by iodine vapor. Each image is a representative of three independent experiments conducted using three independent sample preparations. See Figure S6 for all data. **(B)** Phospholipid composition was determined by densitometry from four consecutive images of each plate, taken every two minutes, and averaging across three independent experiments. The average values are shown as pie charts with the numerical values at the bottom. **(C)** Statistical significance of the difference in PS content shown in B among uninfected and virus-infected Vero ER was analyzed using one-way ANOVA (ns, not significant; \**p* < 0.05; \*\**p* < 0.01) and is presented as the mean ± SD of twelve images, with four images from each of three independent experiments.

## DISCUSSION

Many enveloped viruses, including flaviviruses, exploit PS receptors to gain entry to target cells. However, not all PS receptors can be utilized equally well. For instance, as we previously reported, ZIKV, but not WNV or DENV, can use AXL, as ZIKV is the only one capable of binding GAS6 (31). Because all PS receptors recognize PS in the viral membrane rather than viral proteins, and because the membrane of enveloped viruses is derived from the host cell membrane, the differential use of PS receptors by closely related viruses is puzzling, particularly when those viruses are grown in the same cells. In the current study, we found that the PS content of the virion membrane varies among these viruses, even when produced in the same cells. The higher PS content of ZIKV compared to WNV and DENV explains why only ZIKV can bind GAS6. This finding reveals a previously unrecognized mechanism by which lipid composition of the virion membrane drives receptor usage.

Although there are multiple lipidomic studies on virus-infected cells, only a few have focused on virion lipid composition (71–75). Two studies report PS contents of non-flaviviral virions: while HIV-1 contains higher PS than cellular membranes (71), HCV shows no detectable PS (74). One study on flavivirus found that WNV grown in HeLa cells had higher PS compared to the host cell membrane (75), which contradicts our findings. This discrepancy may, in part, be due to differences in analytical methods: this lipidomic study used mass spectrometry, whereas we used 2D TLC. While mass spectrometry is a powerful tool for many applications, it is less reliable for quantifying complex lipids by class due to variable ionization efficiencies of different molecular species (e.g., those sharing the same head group but differing in fatty acyl chains) and the limited availability of comprehensive internal standards. In contrast, 2D TLC directly separates and visualizes lipids by class without being affected by these factors.

Searching for a potential explanation for why WNV and DENV cannot bind GAS6, we initially investigated virion maturity. In mature flaviviruses, the virion surface is completely covered with E protein homodimers arranged in a herringbone pattern, leaving little membrane exposed (16, 76, 77). However, even in the mature virion, patches of virion membrane are transiently exposed especially at 37°C (19), because virion shells are in a continuous dynamic motion (21, 78), which is described as “breathing” (22). Immature virions, on the other hand, expose larger portions of their membrane because of the standing configuration of prM-E heterotrimers (21). While these immature virions can bind PS receptors, their infectivity remains unclear. Mosaic virions, which are partially immature and partially mature (21, 22, 79, 80), have both immature regions capable of binding PS receptors and mature regions that can mediate infection. Therefore, if the ZIKV population contains a higher proportion of mosaic or immature virions compared to WNV and DENV, this could explain why only ZIKV binds GAS6. However, our data show that WNV engineered to be immature still does not bind GAS6, indicating that virion maturity does not contribute to GAS6 binding. Rather, we found that WNV and DENV have barely detectable levels of PS in their virion, while ZIKV has PS levels comparable to that of the ER membrane of the Vero cells in which the viruses were propagated. Because WNV and DENV have low PS content, even when WNV was made more immature, it could not bind GAS6.

We hypothesized that at least three different scenarios could explain lower PS content in WNV and DENV compared to ZIKV and Vero ER. First, infection of cells by different flaviviruses differentially alter the cellular lipidome, especially in the expanded ER where flaviviruses replicate and assemble. We investigated this possibility and found this was not the case. PL composition of the ER membrane did not notably change following infection by any of the three viruses. We deduce from these results that the viruses themselves or virus replication-assembly processes must be responsible for their distinct virion phospholipid compositions. One possible mechanism for the distinct virion PS content could be the location of virus budding. Flavivirus infection of cells markedly increases synthesis of lipids, including phospholipids, to expand ER membrane, where virus replication takes place. Although all flaviviruses replicate inside the pockets formed on the expanded ER membrane and bud into ER lumen, it is unknown whether all three viruses form replication pockets at similar locations on the expanded ER membrane.

Although PS content of the ER membrane is known to be approximately 4-5% (70, 81, 82), it does not mean PS is evenly distributed in the ER. Rather, different compartments of the ER membrane are believed to have a distinct phospholipid composition. For example, two major enzymes synthesizing PS—Phosphatidyl Synthase 1 (PSS1) that synthesizes PS from PC and Phosphatidyl Synthase 2 (PSS2) that produces PS from PE—are located in the mitochondria-associated membrane (MAM) in mammals (83) and synthesize majority of cellular PS. Much of PS produced in this location is funneled to the mitochondria and converted to PE by Phosphatidylserine Decarboxylase that resides inside the mitochondria (83), and PE produced therein is transported back to ER through MAM and converted to PS and PC. Thus, MAM, a subdomain of ER that connects ER to the nearby mitochondria (84), is enriched with PS and PE. Therefore, although all three viruses bud from the ER membrane, it is possible ZIKV forms replication sites closer to MAM than do DENV or WNV, receiving a higher PS level than do DENV and WNV.

In summary, our data demonstrate that PS content of the closely-related ZIKV, DENV, and WNV viruses are quite different and that higher PS content is linked to ZIKV structural proteins.

## MATERIALS AND METHODS

### Cell lines

HEK293T (human embryonic kidney), Huh-7 (human hepatoma), and Vero (monkey kidney) cells were grown in high-glucose DMEM, and A549 (human lung) and LoVo (human colon) cells were grown in F-12K medium. All cells were cultured in medium supplemented with 10% FBS, at 37°C with 5% CO_2_.

### Virus production

ZIKV Brazilian strain PB-81 was obtained from the World Reference Center for Emerging Viruses and Arboviruses (WRCEVA) at University of Texas Medical Branch (UTMB), DENV type 2 New Guinea C strain was obtained from ATCC (VR-1584), and WNV lineage I New York 1999 (NY99) strain was kindly provided by R. Tesh, UTMB, Galveston, TX. All these flaviviruses were propagated in Vero cells at 37°C in DMEM supplemented with 10% FBS, and the culture supernatants containing viruses were clarified using 0.45 um filters. Flaviviruses were also produced from Huh7, A549, and LoVo cell lines in the respective cell culture medium and filtered. To assess virus titers, viral RNA was extracted and quantified by RT-qPCR as described below, using the primers and probe listed in Table 1. Aliquoted viruses were kept at -80°C. Although not required for ZIKV and DENV, all live virus experiments were conducted side by side with WNV in the BSL3 facility at UF Scripps Institute for Biomedical Research with an approval from Institutional Biosafety Committee.

### Immature virus production

Immature WNV particles were produced by culturing WNV-infected Vero cells in DMEM containing 10% FBS and 50 uM Decanoyl-RVKR-CMK (a furin inhibitor), or by culturing the cells at 28°C without the furin inhibitor, or growing WNV in LoVo cell line that has a mutation in the furin gene (85), Virus was harvested from Vero cells 2d later regardless of culturing temperature and 3d later from LoVo cells. Viruses were quantified by RT-qPCR as described below, and immature status of WNV virions was assessed by detecting uncleaved prM using the antibody against WNV M protein (Abcam, Cat# ab25888).

### Ig-fusion protein production

The plasmids expressing GAS6-Ig, C-GAS6-Ig or TIM1-Ig were described previously (31). Briefly, the cDNA encoding either full-length human GAS6 (GenBank accession no. AAH38984.1), the C-terminal domain of GAS6 (residues 279-678), or the ectodomain of human TIM1 (residues 1-290, GenBank accession no. AAC39862.1) were inserted between the CD5 signal peptide and the Fc domain of human IgG1 that were previously cloned in the pcDNA3.1(+) vector. The plasmid encoding γ-glutamyl carboxylase (GGCX) was described previously (86) and generated by cloning its cDNA sequence into pQCXIP vector.

To produce GAS6-Ig or C-GAS6-Ig, the corresponding expression plasmid was co-transfected with the plasmid encoding GGCX into HEK293T cells at a ratio of 3:1, using calcium-phosphate method. At 6 h post-transfection, cells were washed completely with Phosphate-buffered saline (PBS) to remove remaining FBS and replenished with the FreeStyle 293 medium (Gibco, Cat#12338-018) supplemented with 10 μg/ml vitamin K1 (Sigma). In parallel, TIM1-Ig was produced in FreeStyle 293 medium by transfecting its expression plasmid into HEK293T cells. Mock-transfected culture supernatant was produced by transfecting cells with an empty plasmid and used as a negative control. The culture supernatants containing Ig-fusion proteins or mock-transfected supernatant were harvested at 72 h post-transfection and clarified by 0.45-μm filtration. These Ig-fusion proteins were quantified by SDS-PAGE and Coomassie blue staining after precipitated by Protein A-Sepharose, using known amount of purified human IgG treated in the same way as a standard.

### Virus pull-down assays

2 × 10^8^ genome copies of infectious ZIKV, DENV, or WNV particles were diluted to 400 µl in Tris-buffered saline (TBS) containing 10mM CaCl_2_. Diluted virus was then mixed with 0.2 μg GAS6-Ig, C-Gas6-Ig, or TIM1-Ig protein in 100 µl FreeSytle 293 medium and incubated at 37°C for 1 h. As a negative control for the Ig-fusion proteins, 100 µl of mock-transfected culture supernatant was incubated with the same amount of diluted virus particles. Then, 50 µl Protein A-Sepharose beads, blocked with 2% bovine serum albumin (BSA) and washed, were added to the virus and Ig-fusion protein mixture and incubated for an additional hour at room temperature (RT) with continuous rocking. Captured viruses were analyzed by WB analysis and/or RT-qPCR after Protein A beads were washed three times with 500 µl of TBS containing 0.05% Tween-20 (vol/vol) (TBST).

### Virus WB

Captured viruses were analyzed by both nonreducing (to detect viral E proteins) and reducing (to detect Ig-fusion proteins) SDS-PAGE. For E protein blotting, proteins on the non-reducing gels were transferred to polyvinylidene difluoride (PVDF) membranes and blotted with either 4G2 antibody, which recognizes ZIKV and DENV E protein, or 3.91D antibody (Millipore, Cat# MAB8151), which binds WNV E protein. The 4G2 antibody was produced and purified from the Hybridoma cell line (ATCC, Cat# HB-112). To assess immaturity of WNV, anti-WNV M protein (Abcam, Cat# ab25888) was used. For Ig-fusion protein detection, reducing gels were transferred to PVDF membrane and blotted with a horseradish peroxidase (HRP)-conjugated goat-anti-human Fcγ (Jackson Immuno Research Laboratories, Cat# 109-035-098).

### Virus RT-qPCR

To titer produced viruses and to quantify captured viruses in the pull-down assays, viral RNA was extracted either from the cell culture supernatant or the precipitated beads, using TRIzol (Invitrogen, Cat# 10296028) and GlycoBlue coprecipitant (Invitrogen, Cat# AM9516). Extracted RNA was reverse transcribed using a high-capacity cDNA reverse transcription kit (Applied Biosystems, Cat# 4374966). qPCR was performed using Luna Universal Probe qPCR Master Mix (New England Biolabs, Cat# M3004) with the specific primers and probes targeting the NS3 gene of ZIKV, DENV, or WNV (Table 1), using the PCR protocol: 95°C for 3min x 1 cycle, followed by 95°C for 5 sec and 60°C for 30 sec x 40 cycles. Known quantity of a plasmid containing the target NS3 gene fragment of each virus was used to generate standard curves.

**Table 1.**
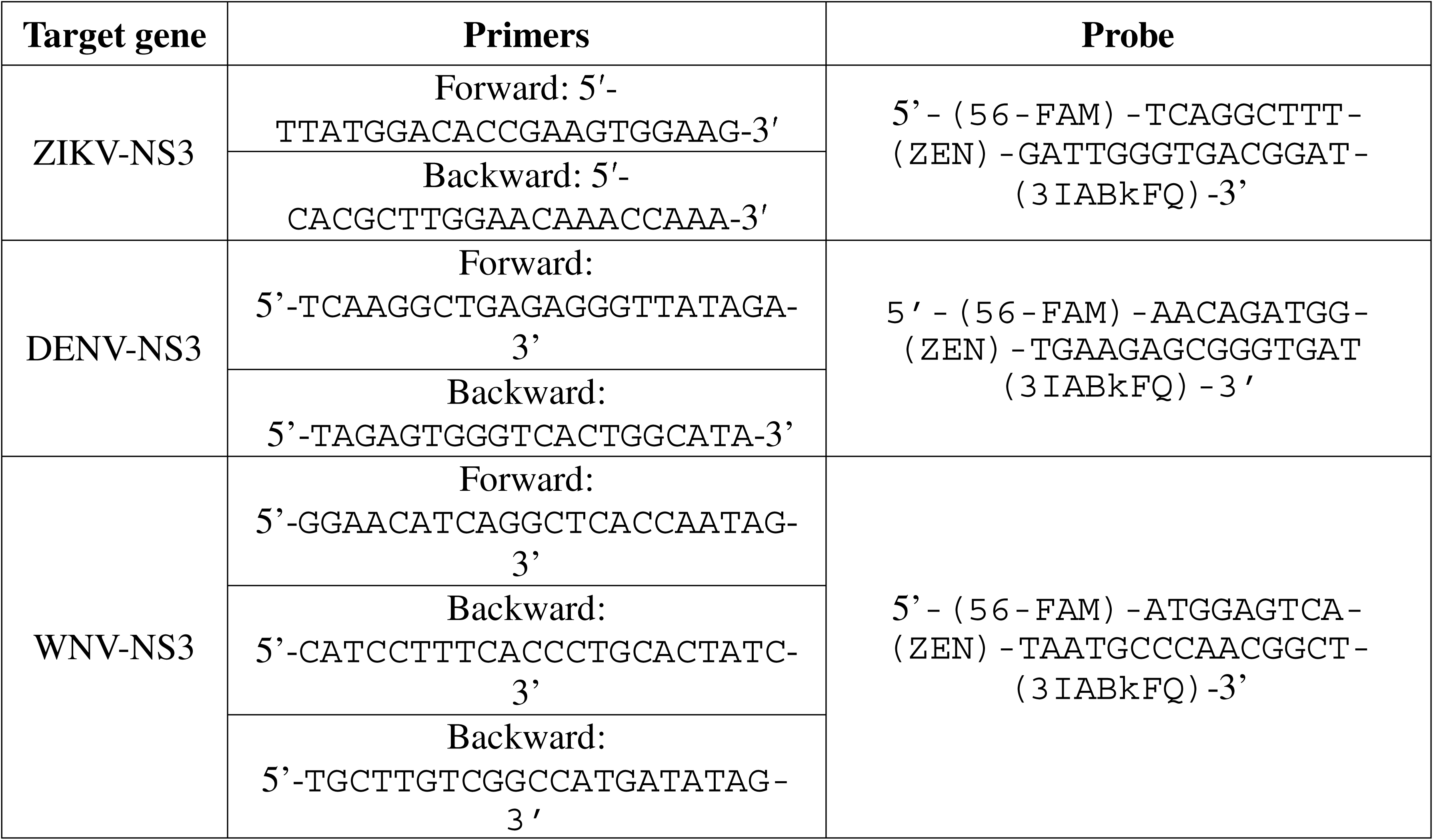
The sequences of qPCR primers and probes used to quantify live viruses.

### Removal of virion N-glycans

WNV particles of 1.0 × 10^9^ genome copies, produced from Vero cells, were treated with 1000 units of PNGase F (NEB, Cat# P0704) in 50 ul medium containing 1X GlycoBuffer 2 at 37°C for 4 h, and the removal of N-glycans was confirmed by reduction in apparent size of the E protein by WB.

### Phospholipase C treatment of virion

ZIKV grown in Vero cells (approximately 4 × 10^8^ genome copies in 200 ul) was treated with phospholipase C (PLC) at 1 unit/ml at 37°C for 1h.

### Phospholipid ELISA

Synthetic phospholipids, 12-dioleoyl-sn-glycero-3-phosphocholine (DOPC), 12-dioleoyl-sn-glycero-3-phosphoethanolamine (DOPE), 12-dioleoyl-sn-glycero-3-[phospho-l-serine] (DOPS), or 1,2-dioleoyl-sn-glycero-3-phospho-[1’-myo-inositol] (DOPI) were purchased from Avanti Polar Lipids. Polystyrene ELISA plates were coated with the indicated amount of these lipids and air dried completely over night as described previously (86). The plates were then blocked with 1% BSA in TBS, incubated with 3 nM GAS6-Ig or TIM1-Ig in TBS containing 10 mM CaCl_2_ (TBS-Ca^2+^) at RT for 1 h, washed with TBST-Ca^2+^ containing 0.05% (vol/vol) Tween 20, and incubated with an HRP-conjugated goat-anti-human Fcγ (Jackson Immuno Research Laboratories, Cat# 109-035-098). After three washes with TBST-Ca^2+^, binding was visualized using UltraTMB substrate (Thermo Scientific Pierce, Cat# 34028). The reaction was terminated with 2 M phosphoric acid, and the plates were read at 450 nm with a SpectraMax Paradigm microplate reader (Molecular Devices).

### Virus purification

Vero cells were infected with ZIKV, DENV, or WNV at an MOI of 1, and cultured in high glucose DMEM supplemented with 10% FBS. Cell culture supernatants containing viruses were collected 2 days later (ZIKV and WNV) or 6 days later (DENV), when 20-30% of the cells showed cytopathic effect. The supernatants were then cleared by centrifugation at 500 ×g for 5 minutes, followed by filtration through a 0.45 µm filter. The supernatants were supplemented with polyethylene glycol 8000 (PEG-8000) to 8% (w/v), incubated with rocking at 4°C overnight, and precipitated by centrifugation at 3,000 ×g for 1 h at 4°C the next day. Pelleted viruses were then resuspended in 10 ml NTE buffer (120 mM NaCl, 20 mM Tris-HCl, 1 mM EDTA, pH 8.0), loaded onto a 2 ml 24% sucrose cushion, and centrifuged in an SW41 rotor at 175,000 ×g for 2 hours at 10°C. Pelleted viruses were resuspended in 1 ml NTE buffer and subjected to a linear 10–35% (w/v) potassium tartrate gradient centrifugation at 175,000 ×g for 2 hours at 10°C. The virus band, located approximately at 20% in the potassium tartrate gradient, was harvested and pelleted by centrifugation at 175,000 ×g for 2 hours at 10°C. Purified viruses were resuspended in NT buffer (120 mM NaCl, 20 mM Tris-HCl, pH 8.0) and stored at -80°C.

### Endoplasmic reticulum (ER) isolation

Total ER (both rough and smooth ER) was isolated from the mock- and virus-infected Vero cells using the ER Enrichment Extraction kit (Novus Biologicals, Cat# NBP2-29482) by following the manufacture’s instruction. Briefly, three T175 flasks of over 90% confluent uninfected Vero cells or those infected with viruses were harvested at day 2 post WNV and ZIKV infection or day 6 post DENV infection. Cells were washed once with PBS, pelleted at 500 × g for 5 minutes, and then resuspended in 1X isosmotic homogenization buffer (IHB) supplemented with 1X protease inhibitor cocktail (PIC) provided by the kit. These cells were homogenized on ice in a clean PTFE tissue grinder (Cole-Parmer, Cat# UX-44468-02). Homogenates were centrifuged at 1,000 ×g for 10 min at 4°C to eliminate pelleted nuclei and cell debris, followed by another centrifugation at 12,000 ×g for 15 min at 4°C to further remove cell debris and mitochondria. Then, supernatants were transferred to ultracentrifuge tubes, centrifuged at 130,000 ×g for 1 h at 4°C, and resuspended in 200 μl suspension buffer in the kit containing 1X PIC from the kit.

### Lipid extraction from the ER membrane or purified viruses

Total lipids were extracted from purified flavivirus particles or isolated ER using the Bligh & Dyer method (87). For comparing virion lipids, lipids were extracted from 1 × 10¹³ genome copies of ZIKV, DENV, or WNV particles. For ER lipids, isolated ER equivalent of 1 mg of total protein was used for lipid extraction. Specifically, 1.5 ml of methanol and chloroform mixed at a 2:1 ratio was added to a viral or ER sample diluted in 0.4 ml PBS in a glass vial. The mixture was vortexed briefly and allowed to sit for 5 minutes at room temperature, 0.5 ml of chloroform was added to it, and the solution was vortexed for 30 seconds. 0.5 ml of double-distilled H O (ddH O) was then added, and the solution was vortexed again before centrifuged at 250 ×g in a GH-3.8 rotor for 5 minutes at room temperature. The bottom chloroform phase containing the lipids was carefully collected with a Pasteur pipette and transferred to a new glass vial. An equal volume of a 2:2:1 mixture of methanol, chloroform, and ddH O was added and the extraction procedure was repeated. After centrifugation, the bottom phase was collected into a 1.5 ml Eppendorf tube and dried overnight in a fume hood.

### Two-dimensional thin-layer chromatography (2D TLC)

For phospholipid analysis by 2D-TLC, dried viral or ER lipids were resuspended in 100 µl chloroform and loaded onto a TLC plate coated with silica gel 60 F_254_ (0.5 mm, 20 × 20 cm, Millipore, Cat# 1.05744.001) at a spot 2 cm away from the either edge. For a sharper separation, a small amount of sample was loaded and dried completely before another aliquot was loaded. To verify the position of each lipid, a mixture containing synthetic DOPC, DOPE, DOPS, DOPI, and sphingomyelin (SPH) was separated by 2D-TLC in the same way. For 1^st^ dimensional separation, the plate with loaded sample was placed in a sealed TLC chamber for 90 min with a solvent composed of chloroform, methanol, ammonium hydroxide, and water at a ratio of 13:5:1:19. The plate was air-dried for 30 min and run for 2^nd^ dimension for 90 min in a solvent containing chloroform, acetone, methanol, acetic acid, and water at a ratio of 10:4:2:2:1. After drying, the plate was stained with iodine vapor produced from iodine crystals heated at 40°C in a sealed chamber placed inside a fume hood. The plate was removed from the iodine chamber, placed in a fume hood, and let iodine to evaporate until background yellow color dissipated and phospholipid spots became clear. During this process, images were captured every two minutes in a ChemiDoc imager (Bio-Rad), and the intensity of phospholipid spots was analyzed using the Image Lab software (Bio-Rad). Several images of varying intensity were analyzed and averaged per plate.

### Statistical analysis

All data were analyzed with GraphPad Prism version 9.0 (GraphPad Software Inc.) and expressed as Mean ± standard error (SD). Statistical significance among the groups was analyzed by one-way ANOVA with Tukey’s multiple comparisons test or two-way ANOVA (ns, not significant; \**P* < 0.05, \*\**P* < 0.01, \*\*\**P* < 0.005, and \*\*\*\**P* < 0.005).

## Supporting information

Figure S1. Purified ZIKV, WNV, and DENV used for lipid analysis.

Figure S2. PS content of ZIKV is substantially higher than that of WNV and DENV.

Figure S3. PL composition of the ER membrane is not differentially altered by different viruses.

## Data, materials, and software availability

All data are included in the main manuscript and supplemental figures.

## AUTHOR CONTRIBUTIONS

B.S., L.Z., and H.C. designed the study. B.S., L.Z., C.K., and Y.K. performed experiments. A.S.R. and M.R.F. contributed key reagents. B.S. and L.Z. analyzed the data. L.Z. and H.C. wrote the manuscript. All authors reviewed and edited the manuscript.

## ACKNOWLEDGEMENTS

We thank Dr. Robert Tesh at University of Texas Medical Branch at Galveston, TX, and the World Reference Center for Arboviruses and Emerging Viruses for providing us with WNV and ZIKV. This work was supported by National Institute of Health grant R01 AI110692 (H.C.)

