## Supplementary figures and images for "Variation in virion phosphatidylserine content drives differential GAS6 binding among closely related flaviviruses"

### Figure S1. Purified ZIKV, WNV, and DENV used for lipid analysis.

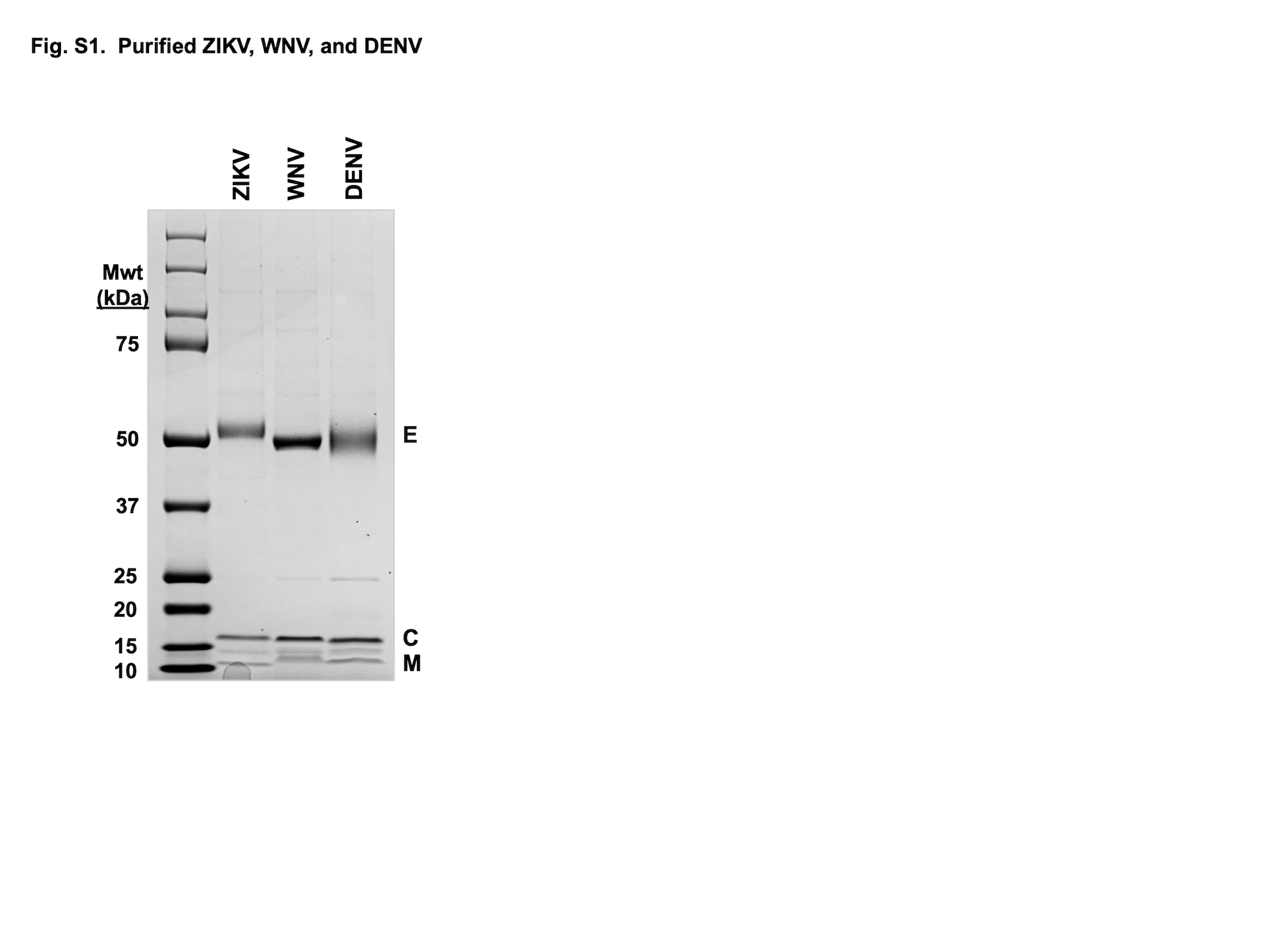

### Figure S2. PS content of ZIKV is substantially higher than that of WNV and DENV.

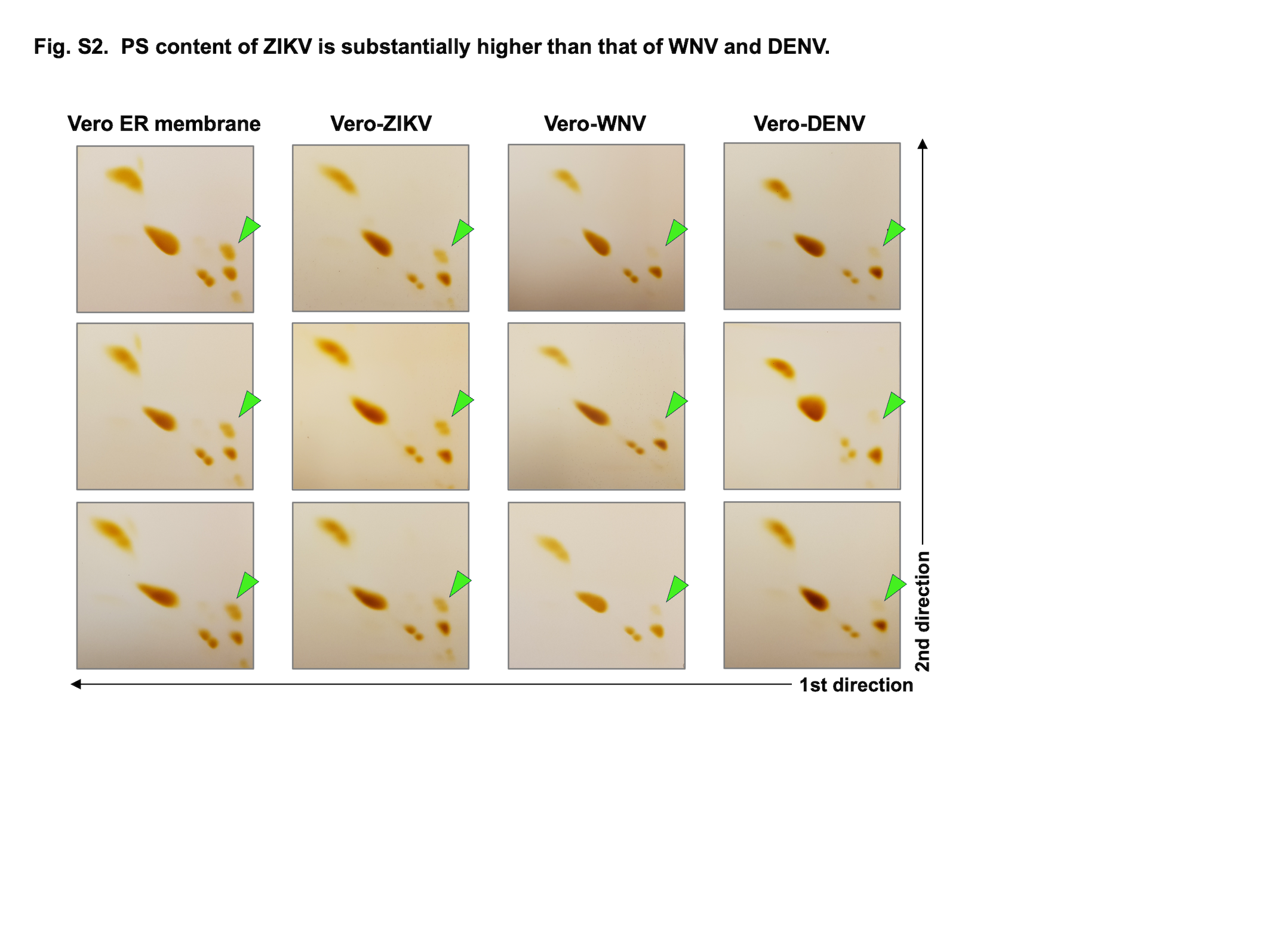

### Figure S3. PL composition of the ER membrane is not differentially altered by different viruses.

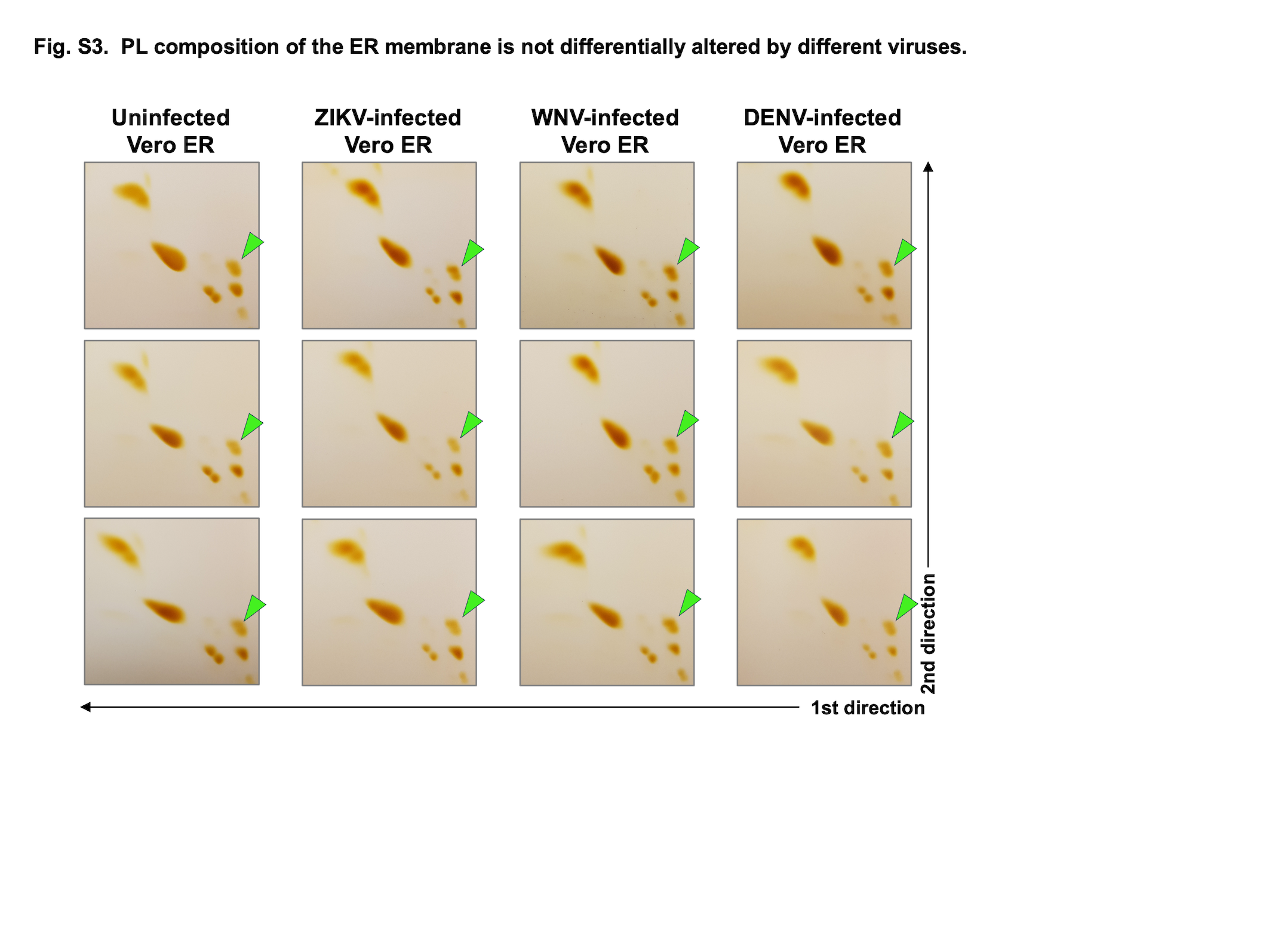
